# Improving ADMET prediction with descriptor augmentation of Mol2Vec embeddings

**DOI:** 10.1101/2025.07.14.664363

**Authors:** Roman Stratiichuk, Nazar Shevchuk, Roman Kyrylenko, Volodymyr Vozniak, Ihor Koleiev, Taras Voitsitskyi, Sasha Korogodski, Zakhar Ostrovsky, Ivan Khropachov, Valerii Vasylevskyi, Vladyslav Husak, Serhii Starosyla, Semen Yesylevskyy, Alan Nafiiev

## Abstract

The accurate prediction of ADMET (Absorption, Distribution, Metabolism, Excretion, and Toxicity) properties is crucial for early-stage drug development, enabling the reduction of late-stage attrition and guiding compound prioritization. In recent years, machine learning models have emerged as powerful tools for ADMET prediction, leveraging diverse molecular representations ranging from handcrafted descriptors to graph neural networks and language model embeddings. Despite these advances, balancing predictive performance with computational efficiency remains a key challenge, particularly for high-throughput screening scenarios. Among unsupervised embedding methods, Mol2Vec has shown promise by capturing chemical substructure context analogously to word embeddings in natural language processing. However, its performance on comprehensive ADMET benchmarks has not been systematically assessed. In this work, we reimplement Mol2Vec with an expanded training corpus and higher embedding dimensionality, and evaluate its utility across 16 ADMET prediction tasks from the Therapeutics Data Commons (TDC). We show that while Mol2Vec embeddings alone are competitive, combining them with classical molecular descriptors and applying feature selection significantly improves performance. Our final MLP models with enhanced Mol2Vec embeddings achieved top-1 results in 10 of 16 benchmarks, outperforming all previously reported models on the TDC leaderboard in this regard, demonstrating that descriptor-enriched representations, paired even with relatively simple MLPs, can rival or exceed the performance of more complex models.

## 1. Introduction

Achieving favorable pharmacokinetic and toxicity profiles, collectively referred to as ADMET (Absorption, Distribution, Metabolism, Excretion, and Toxicity), remains one of the principal challenges in modern drug development. Experimental ADMET assays, while reliable, are costly and time-consuming, making them less feasible for high-throughput screening in the early phases of drug discovery. As a result, *in silico* methods, particularly those based on Quantitative Structure–Activity/Property Relationships (QSAR/QSPR), have become essential tools for the preliminary evaluation of drug-like candidates [1].

Recent advances in machine learning (ML) have led to widespread adoption of data-driven models for ADMET prediction. Trained on extensive experimental datasets, these models can estimate pharmacokinetic and toxicity properties of compounds prior to synthesis, thereby accelerating the design–make–test cycle. Given the increasing reliance on ML models, rigorous benchmarking and transparent performance evaluation are critical for assessing their real-world applicability.

The Therapeutics Data Commons (TDC) platform addresses this need by providing standardised datasets, unified data processing tools, and an active leaderboard for comparative evaluation of ADMET prediction models [2]. TDC currently includes 22 curated datasets relevant to drug safety and pharmacokinetics, enabling systematic comparison across a broad range of modelling approaches.

A key determinant of model performance in ADMET tasks is the molecular representation used as input. Traditional approaches rely on hand-crafted descriptors such as Extended Connectivity Fingerprints (ECFP) and Mordred descriptors [3,4], which encode molecular features based on predefined structural rules. More recently, large language models have been employed to learn molecular embeddings directly from SMILES strings, as exemplified by ChemGPT in ZairaChem [5,6].

Given the graph-like nature of molecular structures, Graph Neural Networks (GNNs) have also gained popularity [7]. These models learn molecular representations by aggregating information across atomic graphs, capturing intricate topological and chemical relationships. Leading models on the TDC leaderboard leverage GNN-based encoders for end-to-end learning. However, recent findings suggest that GNN-derived embeddings can be further improved by incorporating classical descriptors. A notable example is MapLight, which integrates multiple descriptor sets, including ECFP, Avalon [8], ErG fingerprints [9], and over 200 molecular properties, with GNN-based embeddings to achieve state-of-the-art results [10,11].

Another competitive approach is DeepPurpose, a flexible deep learning framework for drug– target interaction prediction [12]. Among its various configurations, models using Morgan or RDKit2D fingerprints with a multilayer perceptron (MLP) encoder perform particularly well on TDC ADMET benchmarks and serve as strong baselines in our comparative study.

These examples illustrate that combining diverse descriptor types, both learned and handcrafted, can significantly enhance model performance. However, the large number of features can pose challenges for model training, especially in small datasets where overfitting is a concern. To address this, feature selection techniques are commonly applied, such as variance thresholding, correlation filtering, and selection based on feature importance from Random Forest models [13]. We decided to explore the application of Mol2Vec, an unsupervised embedding technique originally inspired by Word2Vec, and enrich this embedding with additional chemically relevant features [14,15]. Mol2Vec represents molecules as continuous vectors by learning substructure embeddings in analogy to words in natural language, effectively capturing chemical context while mitigating issues such as high dimensionality and bit collisions inherent in traditional fingerprints [16].

In this study, we reimplemented Mol2Vec with an expanded corpus and increased embedding dimensionality (from 300 to 512) to better capture molecular diversity. We then evaluated the resulting embeddings on the 16 most relevant and widely used ADMET tasks from the TDC benchmark suite, using combinations of Mol2Vec with various descriptor types (e.g., physicochemical properties, ECFP, Mordred, Avalon). Through systematic feature selection and model optimisation, we demonstrate that the Mol2Vec embeddings with additional descriptors and automated feature selection achieve top ranking in 10 out of 16 tested benchmarks, the absolute best among all TCD techniques, highlighting its utility as an efficient representation for ADMET modelling. The general scheme is displayed in Figure 1.

**Figure 1.**
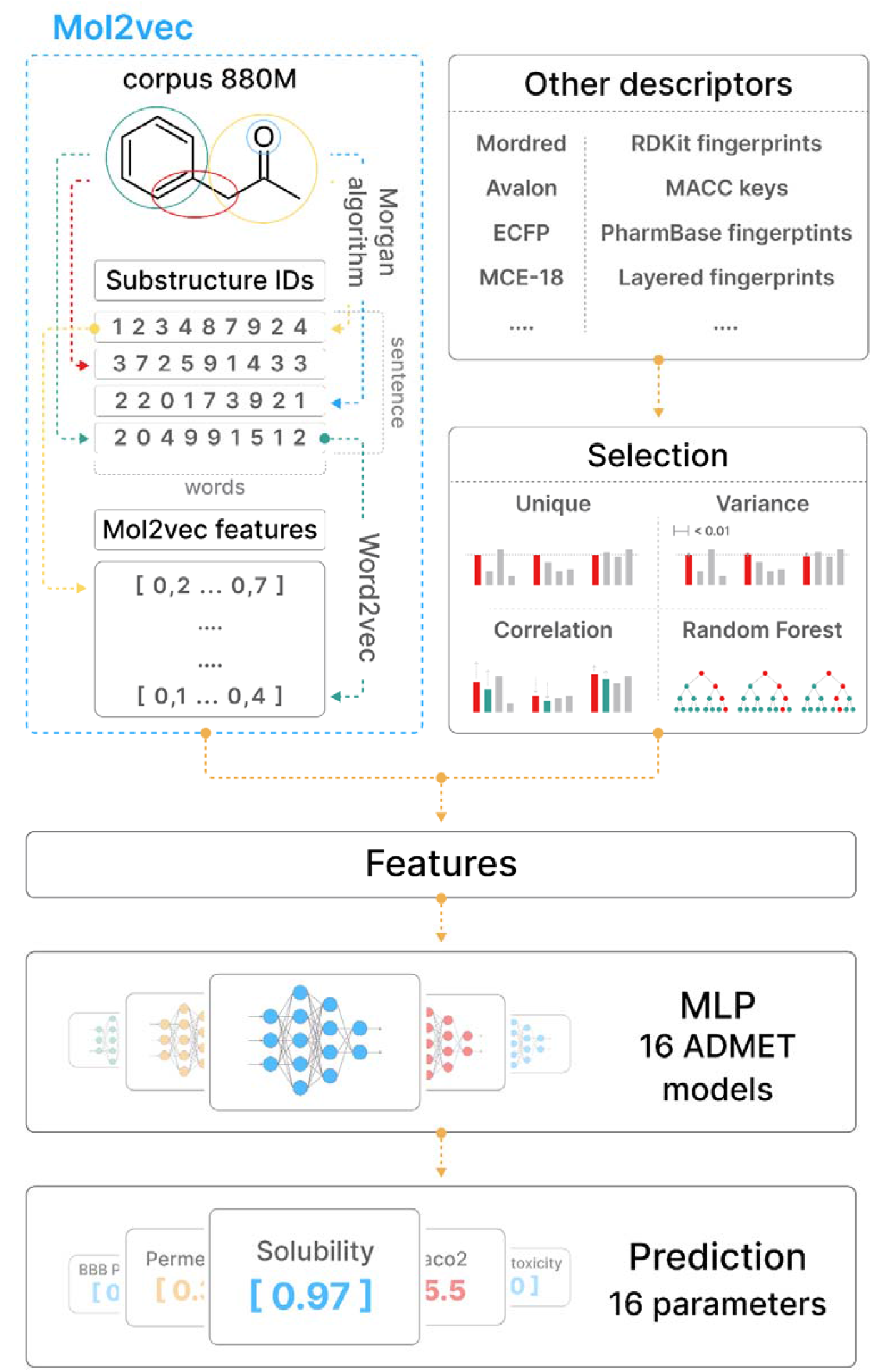
General Workflow for ADMET Prediction Using Mol2Vec Descriptor Augmentation. The pipeline begins with the extraction of molecular substructures using the Morgan algorithm applied to an 880-million compound corpus from ZINC20, generating sequences of substructure IDs (analogous to “sentences”) that are used to train Mol2Vec embeddings via a Word2Vec Skip-gram model. These Mol2Vec embeddings are then combined with various traditional molecular descriptors to form a comprehensive feature set. A feature selection module (using uniqueness filtering, variance thresholding, correlation pruning, and Random Forest importance scoring) reduces this high-dimensional input space. The selected features are then fed into 16 task-specific multilayer perceptron (MLP) models to predict key ADMET properties on 16 TDC benchmarks.

## 2. Materials and Methods

Mol2Vec is an unsupervised method for generating vector embeddings of molecular substructures, which can subsequently be used as input features for supervised learning tasks, such as ADMET prediction in our case. This section describes our Mol2Vec reimplementation, including corpus preparation and model training. We also detail the training and evaluation of ADMET models, focusing on how Mol2Vec embeddings were combined with additional molecular descriptors, the ADMET benchmark datasets used, and the overall training procedure.

### 2.1. Corpus preparation

To construct a robust corpus for Mol2Vec training, we leveraged the ZINC20 database [17], applying a series of filters to ensure chemical relevance and data quality. Specifically, compounds were retained if they possessed a “2D representation,” exhibited “Standard reactivity,” and were flagged as “Wait OK” in terms of commercial availability. This filtering process yielded a dataset comprising 883,897,271 small molecules.

Subsequent preprocessing steps included the removal of counterions and solvent fragments, followed by the generation of canonical SMILES strings using RDKit [18], ensuring consistency and uniqueness in molecular representation. Molecular substructures were extracted using the Morgan algorithm [4] with two radial configurations: radius 0 (capturing atomic identity) and radius 1 (incorporating immediate atomic neighbourhoods). These substructures were treated as “words”, and individual molecules were represented as “sentences” comprising sequences of such words.

The resulting corpus encompassed a vocabulary of 13,278 unique molecular “words.” To mitigate overfitting and enhance generalisation during embedding training, all substructure tokens appearing fewer than 4 times across the entire corpus were replaced with a special “UNK” token, which is in line with standard ML practices for rare token handling.

### 2.2. Mol2Vec embeddings generation

To generate molecular embeddings, we employed two established architectures adapted from natural language processing: Continuous Bag of Words (CBOW) and Skip-gram [19]. The CBOW approach aims to predict a target token based on its surrounding context, whereas Skip-gram performs the inverse operation, predicting contextual tokens from a given target.

We initially evaluated both Continuous Bag of Words (CBOW) and Skip-gram architectures for generating Mol2Vec embeddings. But during the experiments, we found that the Skip-gram model, configured with a context window size of 10 and a minimum substructure occurrence threshold (unseen cut-off) of 4, yielded the most effective molecular representations. This configuration was therefore adopted for all subsequent experiments and final task-specific models training.

Molecular substructure tokens, derived from the corpus as described above, were encoded into 512-dimensional vectors using the Gensim library [20]. Each molecule was represented as a single vector by aggregating the embeddings of all its constituent substructures.

### 2.3. ADMET benchmark datasets

In this study, we selected a subset of 16 benchmarks from the Therapeutics Data Commons (TDC) that are highly relevant to real-world drug discovery applications. These include:

1. 4 benchmarks evaluating drug absorption: Caco-2 cell permeability [21], Human Intestinal Absorption (HIA) [22], P-glycoprotein (Pgp) inhibition [23], and Oral bioavailability [24];
2. 3 benchmarks assessing distribution properties, which describe how a drug disperses throughout the body: Blood-Brain Barrier (BBB) penetration [25,26], Plasma Protein Binding Rate (PPBR) [27], and Volume of Distribution at steady state (VDss) [28];
3. 6 metabolism-related benchmarks, which evaluate how compounds are metabolised and how this affects drug exposure and efficacy: CYP2D6, CYP3A4, and CYP2C9 inhibition, as well as CYP2D6, CYP3A4, and CYP2C9 substrate classification [29];
4. 3 toxicity benchmarks, which assess potential adverse effects: Ames mutagenicity [30], Drug-Induced Liver Injury (DILI) [31], and hERG channel inhibition [32].

Among these tasks, 3 are formulated as regression problems and 13 as binary classification tasks. The dataset sizes vary from small (e.g., HIA with 578 compounds) to large-scale (e.g., CYP2D6 inhibition with 13,130 compounds).

### 2.4. Featurization

To generate input features for our ADMET prediction models, we evaluated several featurization strategies:

1. Mol2Vec only – serving as a baseline representation derived from unsupervised learning of molecular substructures;
2. Mol2Vec embeddings combined with physico-chemical properties – including molecular weight, logP, number of hydrogen bond donors/acceptors, number of rotatable bonds, total polar surface area, formal charge, and other descriptors listed in the Supplementary Table 1;
3. Mol2Vec embeddings with 2D Mordred descriptors – a comprehensive set of hand-crafted molecular features capturing structural and physicochemical properties;
4. Mol2Vec embeddings enriched with a wide range of additional molecular descriptors, followed by automated feature selection procedures – this composite and optimised feature set is referred to as the “Mol2Vec Best” configuration (see Table 1 for descriptor details).

For feature selection in the Mol2Vec Best featurization strategy, we initially considered a set of 94,729 candidate features for each ADMET task. This high-dimensional feature space was systematically reduced using a sequential four-step feature selection pipeline designed to remove redundant, non-informative, or highly correlated features:

**Table 1.** Additional molecular descriptors used in the Mol2Vec Best.

| Descriptors | Number of descriptors | Descriptor parameters |
| --- | --- | --- |
| Mordred descriptors; | 1613 | only 2D |
| MCE-18 descriptor; | 1 | – |
| Morgan fingerprints; | 28672 | bits: [512, 1024, 2048];<br>radius: [1, 2, 3, 4];<br>branches: [True, False] |
| RDKit fingerprints; | 25088 | bits: [512, 1024, 2048];<br>max path: [1, 2, 3, 4, 5, 6, 7] |
| Layered fingerprints; | 25088 | bits: [512, 1024, 2048];<br>max path: [1, 2, 3, 4, 5, 6, 7] |
| Atom fingerprints; | 3584 | bits: [512, 1024, 2048] |
| Topological fingerprints; | 3584 | bits: [512, 1024, 2048] |
| Avalon fingerprints; | 3584 | bits: [512, 1024, 2048] |
| PharmBase fingerprints; | 3584 | – |
| MACCS keys. | 167 | – |

1. Unique selector – features exhibiting a single unique value across all training samples were discarded;
2. Variance selector – features with a variance below 0.05 in the training set were excluded to eliminate those with minimal variability;
3. Correlation selector – eliminated one of any pair of features with a Pearson correlation coefficient exceeding 0.95;
4. Random Forest importance selector – retained features with an importance score greater than 0.9, as computed by a Random Forest Regressor.

These selection steps were applied sequentially to each benchmark dataset. The resulting feature subset was then concatenated with the fixed set of 512 Mol2Vec embedding features to form the final input for model training. The total number of features used for input in each final model is provided in Table 2.

**Table 2.** Total number of features used for input in Mol2Vec Best models after selection.

| Dataset | Selected features | Mol2Vec features | Total |
| --- | --- | --- | --- |
| Caco2 | 1021 | 512 | 1533 |
| HIA | 151 | 512 | 663 |
| Pgp Inhibition | 688 | 512 | 1200 |
| Bioavailability | 1171 | 512 | 1683 |
| BBB | 1329 | 512 | 1841 |
| PPBR | 1997 | 512 | 2509 |
| VDss | 1613 | 512 | 2125 |
| CYP2C9 Inhibition | 6795 | 512 | 7307 |
| CYP2D6 Inhibition | 7588 | 512 | 8100 |
| CYP3A4 Inhibition | 7065 | 512 | 7577 |
| CYP2C9 Substrate | 1124 | 512 | 1636 |
| CYP2D6 Substrate | 7588 | 512 | 2109 |
| CYP3A4 Substrate | 7065 | 512 | 1776 |
| hERG | 794 | 512 | 1306 |
| AMES | 3130 | 512 | 3642 |
| DILI | 639 | 512 | 1151 |

In the following sections, we provide more detailed discussion of ADMET models trained using two specific featurization strategies: Mol2Vec only, which serves as a baseline for evaluating the utility of our reimplemented Mol2Vec approach, and the Mol2Vec Best configuration, which incorporates a broad set of additional molecular descriptors, followed by systematic feature selection. This enhanced configuration consistently achieves superior predictive performance across benchmarks. Results for models trained with other featurization strategies are not discussed, but are summarized in Supplementary Table 3.

### 2.5. ADMET models training

For each ADMET prediction task, a separate Multilayer Perceptron (MLP) model [33] was trained using a task-specific hyperparameter configuration. The model architecture was defined by a fixed set of candidate values for all tunable parameters, which were optimised independently for each dataset.

Hyperparameter optimisation was performed using the Optuna framework v3.3.0 [34], leveraging the Tree-structured Parzen Estimator (TPE) algorithm [35] with standard parameters. The optimisation process was conducted separately for each of the 16 TDC benchmarks, resulting in a distinct set of optimal hyperparameters for each task. Parameters subject to optimisation included:

- Number of hidden layers (ranging from 1 to 5),
- Number of hidden units per layer,
- Dropout rates (from 0.0 to 0.5),
- Activation function (chosen from ELU, ReLU, PReLU, or Leaky ReLU) [36],
- Batch size,
- Weight decay
- Learning rate (lr).

Model training for all benchmarks used the Adam optimizer [37] with the learning rate optimized by Optuna. Each model was trained for 100 epochs, and the number of trials for the tuning process was set to 200. Scaffold-based train/test splits provided by the TDC were used for training. Performance metrics for each endpoint are reported as the mean ± standard deviation, in accordance with TDC evaluation standards.

The full list of optimised hyperparameters for the Mol2Vec Best featurization strategy across all datasets is available in Supplementary Table 2.

## 3. Results and Discussion

Table 3 provides a comparative summary of the current state of the TDC leaderboard, highlighting several representative models that employ distinct molecular featurization strategies. These models were specifically selected for inclusion due to their diversity in encoding approaches and their relevance to our study.

**Table 3.**
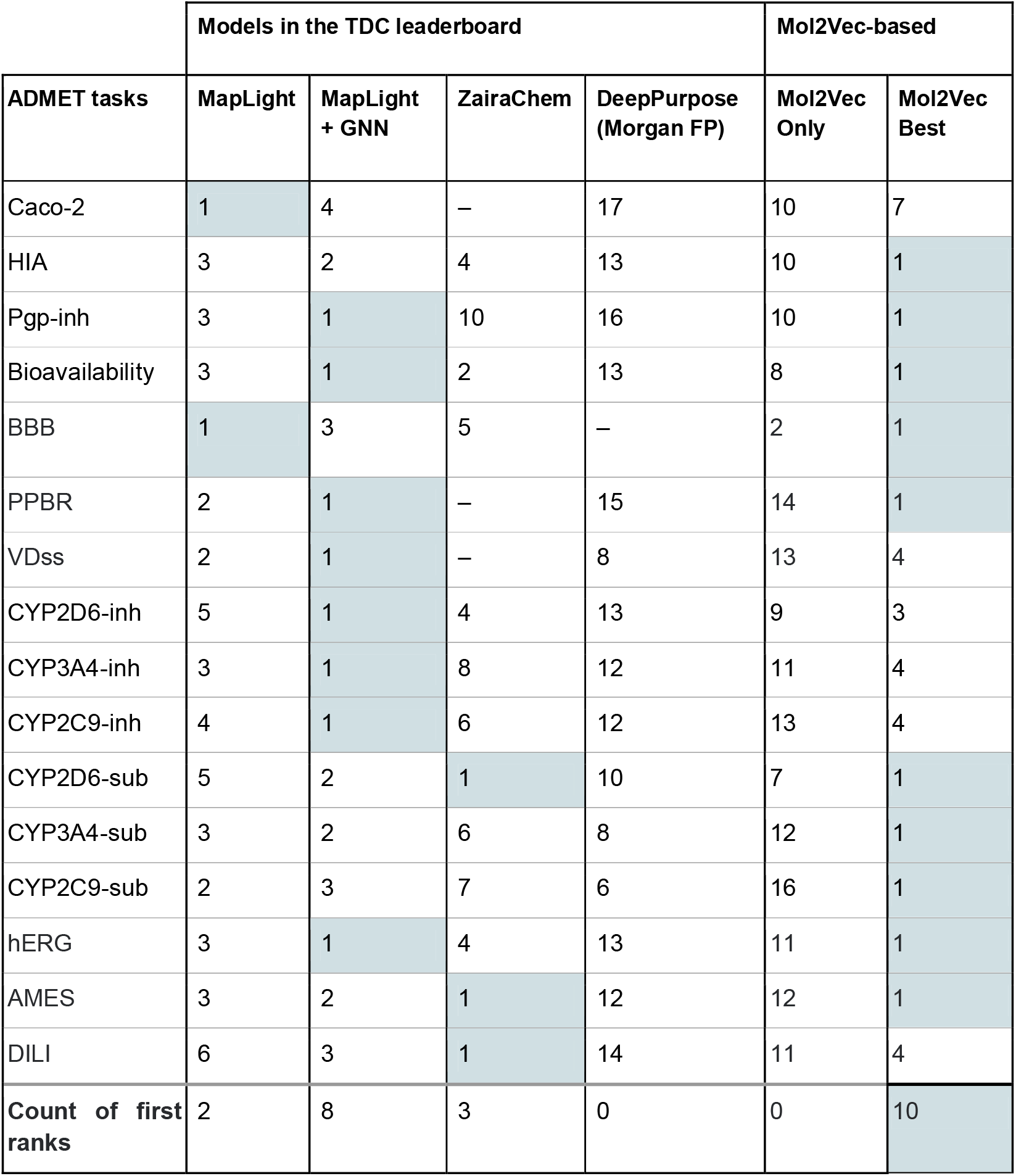
Comparison of TDC leaderboard (as of July 2025) and Mol2Vec-based ADMET models. Model ranks for each ADMET task are reported. The best models for each task are highlighted.

|  | Models in the TDC leaderboard |  |  |  | Mol2Vec-based |  |
| --- | --- | --- | --- | --- | --- | --- |
| ADMET tasks | MapLight | MapLight + GNN | ZairaChem | DeepPurpose (Morgan FP) | Mol2Vec Only | Mol2Vec Best |
| Caco-2 | 1 | 4 | – | 17 | 10 | 7 |
| HIA | 3 | 2 | 4 | 13 | 10 | 1 |
| Pgp-inh | 3 | 1 | 10 | 16 | 10 | 1 |
| Bioavailability | 3 | 1 | 2 | 13 | 8 | 1 |
| BBB | 1 | 3 | 5 | – | 2 | 1 |
| PPBR | 2 | 1 | – | 15 | 14 | 1 |
| VDss | 2 | 1 | – | 8 | 13 | 4 |
| CYP2D6-inh | 5 | 1 | 4 | 13 | 9 | 3 |
| CYP3A4-inh | 3 | 1 | 8 | 12 | 11 | 4 |
| CYP2C9-inh | 4 | 1 | 6 | 12 | 13 | 4 |
| CYP2D6-sub | 5 | 2 | 1 | 10 | 7 | 1 |
| CYP3A4-sub | 3 | 2 | 6 | 8 | 12 | 1 |
| CYP2C9-sub | 2 | 3 | 7 | 6 | 16 | 1 |
| hERG | 3 | 1 | 4 | 13 | 11 | 1 |
| AMES | 3 | 2 | 1 | 12 | 12 | 1 |
| DILI | 6 | 3 | 1 | 14 | 11 | 4 |
| **Count of first ranks** | 2 | 8 | 3 | 0 | 0 | 10 |

In particular, MapLight, a top-performing model across multiple ADMET tasks, utilizes a combination of classical fingerprints and graph-based representations. Its enhanced variant, MapLight + GNN, further integrates graph neural network embeddings and consistently achieves top-1 rankings [38].

ZairaChem employs an ensemble of diverse molecular encoding methods, including language model-based representations, and demonstrates strong performance, although it is limited to classification tasks [6].

DeepPurpose is based on an encoder-decoder architecture that leverages Morgan fingerprints as input to a multilayer perceptron (MLP) encoder, followed by a decoder to predict biological activity. We include DeepPurpose in this comparison due to its architectural similarity to our baseline model employing Mol2Vec embeddings only [12].

Mol2Vec only models show competitive results relative to these benchmarks and surpass the model with the closest architecture – “DeepPurpose Morgan fingerprint + MLP encoder”, validating the utility of unsupervised substructure embeddings for ADMET prediction (Table 3) However, integrating additional molecular descriptors with systematic feature selection applied, as implemented in the Mol2Vec Best configuration, substantially improves predictive performance across most benchmarks. Mol2Vec Best ranks first in 10 out of 16 studied benchmarks, which is the absolute best among all TDC techniques. For Maplight+GNN this value is 8, and the other techniques only show the best ranks in 2-3 benchmarks.

### 3.1. Absorption benchmarks

We achieved top-ranking positions in three out of four absorption-related classification tasks on the TDC leaderboard (Table 4). For all endpoints, excluding Caco-2 permeability, the incorporation of additional molecular descriptors combined with feature selection in the Mol2Vec Best configuration resulted in substantial performance improvements over the Mol2Vec Only baseline. These enhancements were sufficient to elevate our models to first place in the respective benchmark rankings.

**Table 4.** Performance comparison of the TDC top 1 model and Mol2Vec-based models for absorption tasks. Arrows indicate whether higher (↑) or lower (↓) values represent better performance.

| Task | Metric | TDC Best | Mol2Vec Only |  | Mol2vec Best |  |
| --- | --- | --- | --- | --- | --- | --- |
|  |  |  | Value | Rank | Value | Rank |
| Caco-2 | MAE ↓ | $0.276 \pm 0.005$ | $0.377 \pm 0.029$ | 10 | $0.315 \pm 0.017$ | 7 |
| HIA | ROC-AUC ↑ | $0.990 \pm 0.002$ | $0.961 \pm 0.018$ | 10 | $0.996 \pm 0.001$ | 1 |
| Pgp-inh | ROC-AUC ↑ | $0.938 \pm 0.002$ | $0.899 \pm 0.008$ | 10 | $0.948 \pm 0.004$ | 1 |
| Bioavailability | ROC-AUC ↑ | $0.748 \pm 0.006$ | $0.633 \pm 0.027$ | 8 | $0.776 \pm 0.027$ | 1 |

In contrast, the performance gain for the Caco-2 task was minimal, with the Mol2Vec Best model reaching only 7th place. This outcome suggests that the additional descriptors do not provide substantial predictive value for this specific endpoint, possibly due to the limited relevance of the added features or inherent noise in the dataset. Notably, top-performing models in this benchmark category, such as SimGCN [39] and MapLight GNN, utilize graph neural network architectures, indicating that GNN-based encodings may better capture the molecular features relevant to a complex combination of passive and active membrane permeability.

### 3.2. Distribution benchmarks

Our models achieved top-tier performance in two out of the three distribution-related tasks from the TDC benchmark suite (Table 5). Notably, the Mol2Vec-only model attained second place in the Blood-Brain Barrier (BBB) penetration task, despite the absence of additional descriptors. This strong performance indicates that the Mol2Vec embeddings alone effectively capture key structural and physicochemical features relevant to BBB permeability.

**Table 5.**
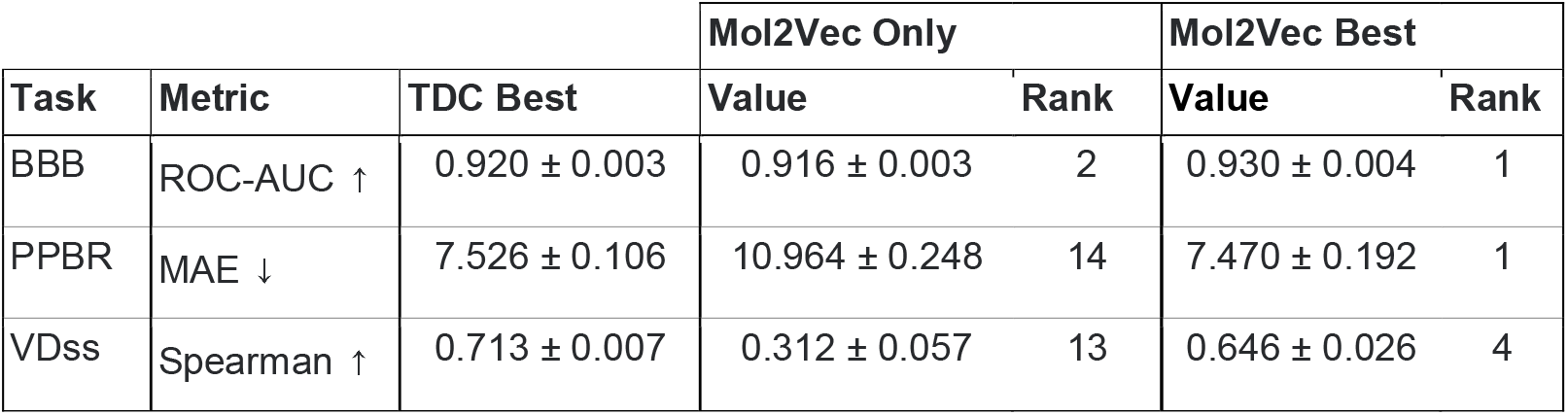
Performance comparison of the TDC top 1 model and Mol2Vec-based models for distribution tasks. Arrows indicate whether higher (↑) or lower (↓) values represent better performance.

| Task | Metric | TDC Best | Mol2Vec Only |  | Mol2Vec Best |  |
| --- | --- | --- | --- | --- | --- | --- |
|  |  |  | Value | Rank | Value | Rank |
| BBB | ROC-AUC ↑ | 0.920 ± 0.003 | 0.916 ± 0.003 | 2 | 0.930 ± 0.004 | 1 |
| PPBR | MAE ↓ | 7.526 ± 0.106 | 10.964 ± 0.248 | 14 | 7.470 ± 0.192 | 1 |
| VDss | Spearman ↑ | 0.713 ± 0.007 | 0.312 ± 0.057 | 13 | 0.646 ± 0.026 | 4 |

For the remaining tasks, the Mol2Vec Best configuration further enhances performance and secures leading positions on the leaderboard.

### 3.3. Metabolism benchmarks

Our Mol2Vec Best models demonstrated superior performance on the majority of cytochrome P450 substrate classification tasks within the TDC benchmark suite (Table 6). While the Mol2Vec Only models underperformed relative to state-of-the-art competitors, the incorporation of additional molecular descriptors in the Mol2Vec Best configuration resulted in substantial performance gains. This enhancement enabled our models to achieve top rankings across multiple CYP-related tasks. Notably, other high-performing models on the TDC leaderboard include MapLight GNN, which excels at CYP inhibition prediction, and ZairaChem, particularly for CYP2C9 substrate classification. Overall, these results underscore the critical role of enriched molecular representations in capturing the complex biochemical determinants of drug metabolism.

**Table 6.** Performance comparison of the TDC top 1 model and Mol2Vec-based models for metabolism tasks. Arrows indicate whether higher (↑) or lower (↓) values represent better performance.

| Task | Metric | TDC Best | Mol2Vec Only |  | Mol2Vec Best |  |
| --- | --- | --- | --- | --- | --- | --- |
|  |  |  | Value | Rank | Value | Rank |
| CYP2D6-inh | PR-AUC ↑ | 0.790 ± 0.001 | 0.632 ± 0.016 | 9 | 0.726 ± 0.004 | 3 |
| CYP3A4-inh | PR-AUC ↑ | 0.916 ± 0.000 | 0.837 ± 0.006 | 11 | 0.884 ± 0.001 | 4 |
| CYP2C9-inh | PR-AUC ↑ | $0.859 \pm 0.001$ | $0.733 \pm 0.011$ | 13 | $0.800 \pm 0.001$ | 4 |
| CYP2D6-sub | PR-AUC ↑ | $0.736 \pm 0.024$ | $0.678 \pm 0.035$ | 7 | $0.821 \pm 0.008$ | 1 |
| CYP3A4-sub | ROC-AUC ↑ | $0.667 \pm 0.019$ | $0.598 \pm 0.042$ | 12 | $0.776 \pm 0.015$ | 1 |
| CYP2C9-sub | PR-AUC ↑ | $0.441 \pm 0.033$ | $0.345 \pm 0.033$ | 16 | $0.556 \pm 0.055$ | 1 |

### 3.4. Toxicity benchmarks

Our Mol2Vec Best models outperformed current state-of-the-art approaches across all three TDC toxicity endpoints: hERG channel inhibition, Ames mutagenicity, and drug-induced liver injury (DILI) (Table 7). In contrast, the Mol2Vec Only models exhibited suboptimal performance, indicating that unsupervised substructure embeddings alone are insufficient to capture the complex biological signals associated with toxicity. The substantial performance gains highlight the importance of integrating diverse molecular descriptors with targeted feature selection. These improvements are particularly evident when compared to strong baseline models such as MapLight GNN (notably effective for hERG) and ZairaChem (which performs well on Ames and DILI), further validating the effectiveness of our composite feature strategy.

**Table 7.** Performance comparison of the TDC top 1 model and Mol2Vec-based models for toxicity tasks. Arrows indicate whether higher (↑) or lower (↓) values represent better performance.

| Task | Metric | TDC Best | Mol2Vec Only |  | Mol2Vec Best |  |
| --- | --- | --- | --- | --- | --- | --- |
|  |  |  | Value | Rank | Value | Rank |
| hERG | ROC-AUC ↑ | $0.880 \pm 0.002$ | $0.753 \pm 0.03$ | 11 | $0.897 \pm 0.003$ | 1 |
| AMES | ROC-AUC ↑ | $0.871 \pm 0.002$ | $0.808 \pm 0.004$ | 12 | $0.876 \pm 0.002$ | 1 |
| DILI | ROC-AUC ↑ | $0.925 \pm 0.005$ | $0.863 \pm 0.034$ | 11 | $0.964 \pm 0.004$ | 1 |

## 4. Conclusion

This study presents a systematic evaluation of Mol2Vec embeddings for ADMET property prediction, demonstrating that lightweight, unsupervised molecular representations can achieve strong performance when combined with classical descriptors and rigorous feature selection. By retraining Mol2Vec on an expanded chemical corpus and increasing the embedding dimensionality to 512, we significantly improved its capacity to capture structural and physicochemical information. While Mol2Vec alone provided competitive results, outperforming certain established baselines such as DeepPurpose, its integration with descriptor sets like Mordred, Avalon, and fingerprint-based encodings yielded consistent and substantial improvements across nearly all tasks.

Our Mol2Vec Best configuration ranked first on 10 of the 16 ADMET benchmarks from the Therapeutics Data Commons and placed within the top five for nearly all others. This constitutes the highest number of top-1 benchmark results among all models reported on the TDC leaderboard. This demonstrates that descriptor-enriched embeddings, optimized through principled feature selection, can rival or even outperform more complex architectures such as GNNs and large language model-based approaches, as shown by our Mol2Vec best embedding.

## 5. CRediT authorship contribution statement

**Roman Stratiichuk:** Conceptualization, Methodology, Project administration, Data curation, Validation, Supervision, Writing – original draft. **Nazar Shevchuk:** Writing – original draft. **Roman Kyrylenko:** Writing – original draft, Writing – review & editing. **Volodymyr Vozniak:** Software, Formal analysis, Investigation, Data curation, Visualization. **Ihor Koleiev:** Methodology, Resources, Investigation, Data curation. **Taras Voitsitskyi:** Resources, Software. **Zakhar Ostrovsky:** Resources, Software. **Ivan Khropachov:** Resources, Software. **Valerii Vasylevskyi:** Data curation, Validation. **Sasha Korogodski:** Data curation, Validation. **Vladyslav Husak:** Validation, Data curation. **Serhii Starosyla:** Methodology, Project administration, Supervision. **Semen Yesylevskyy:** Methodology, Project administration, Supervision, Funding acquisition, Writing – review & editing. **Alan Nafiiev:** Conceptualization, Methodology, Project administration, Supervision.

## 7. Acknowledgements

Semen Yesylevskyy has received funding through the grant MSMT-355/2025-16 from the Ministry of Education, Youth and Sport of the Czech Republic and through the MSCA4Ukraine project 101101923, which is funded by the European Union. Views and opinions expressed are, however, those of the author(s) only and do not necessarily reflect those of the European Union. Neither the European Union nor the MSCA4Ukraine Consortium as a whole nor any individual member institutions of the MSCA4Ukraine Consortium can be held responsible for them. The authors thank Prof. Alexander Zholos for his valuable consultations during the preparation of this manuscript.

## References

[1] Wu F, Zhou Y, Li L, Shen X, Chen G, Wang X, et al. Computational Approaches in Preclinical Studies on Drug Discovery and Development. Front Chem 2020;Volume 8-2020.

[2] Huang K, Fu T, Gao W, Zhao Y, Roohani Y, Leskovec J, et al. Therapeutics Data Commons: Machine Learning Datasets and Tasks for Drug Discovery and Development 2021. 10.48550/arXiv.2102.09548.

[3] Moriwaki H, Tian Y-S, Kawashita N, Takagi T. Mordred: a molecular descriptor calculator. J Cheminformatics 2018;10:4. 10.1186/s13321-018-0258-y.

[4] Rogers D, Hahn M. Extended-Connectivity Fingerprints. J Chem Inf Model 2010;50:742–54. 10.1021/ci100050t.

[5] Sadeghi S, Bui A, Forooghi A, Lu J, Ngom A. Can large language models understand molecules? BMC Bioinformatics 2024;25:225. 10.1186/s12859-024-05847-x.

[6] Turon G, Hlozek J, Woodland JG, Kumar A, Chibale K, Duran-Frigola M. First fully-automated AI/ML virtual screening cascade implemented at a drug discovery centre in Africa. Nat Commun 2023;14:5736. 10.1038/s41467-023-41512-2.

[7] Khemani B, Patil S, Kotecha K, Tanwar S. A review of graph neural networks: concepts, architectures, techniques, challenges, datasets, applications, and future directions. J Big Data 2024;11:18. 10.1186/s40537-023-00876-4.

[8] Gedeck P, Rohde B, Bartels C. QSAR − How Good Is It in Practice? Comparison of Descriptor Sets on an Unbiased Cross Section of Corporate Data Sets. J Chem Inf Model 2006;46:1924–36. 10.1021/ci050413p.

[9] Stiefl N, Watson IA, Baumann K, Zaliani A. ErG: 2D Pharmacophore Descriptions for Scaffold Hopping. J Chem Inf Model 2006;46:208–20. 10.1021/ci050457y.

[10] Heid E, Greenman KP, Chung Y, Li S-C, Graff DE, Vermeire FH, et al. Chemprop: A Machine Learning Package for Chemical Property Prediction. J Chem Inf Model 2024;64:9–17. 10.1021/acs.jcim.3c01250.

[11] Notwell JH, Wood MW. ADMET property prediction through combinations of molecular fingerprints 2023. 10.48550/arXiv.2310.00174.

[12] Huang K, Fu T, Glass LM, Zitnik M, Xiao C, Sun J. DeepPurpose: a deep learning library for drug-target interaction prediction. Bioinforma Oxf Engl 2021;36:5545–7. 10.1093/bioinformatics/btaa1005.

[13] Pudjihartono N, Fadason T, Kempa-Liehr AW, O’Sullivan JM. A Review of Feature Selection Methods for Machine Learning-Based Disease Risk Prediction. Front Bioinforma 2022;Volume 2-2022.

[14] Jaeger S, Fulle S, Turk S. Mol2vec: Unsupervised Machine Learning Approach with Chemical Intuition. J Chem Inf Model 2018;58:27–35. 10.1021/acs.jcim.7b00616.

[15] Mikolov T, Chen K, Corrado G, Dean J. Efficient Estimation of Word Representations in Vector Space 2013. 10.48550/arXiv.1301.3781.

[16] Shibayama S, Marcou G, Horvath D, Baskin II, Funatsu K, Varnek A. Application of the mol2vec Technology to Large-size Data Visualization and Analysis. Mol Inform 2020;39:1900170. 10.1002/minf.201900170.

[17] Irwin JJ, Tang KG, Young J, Dandarchuluun C, Wong BR, Khurelbaatar M, et al. ZINC20—A Free Ultralarge-Scale Chemical Database for Ligand Discovery. J Chem Inf Model 2020;60:6065–73. 10.1021/acs.jcim.0c00675.

[18] Landrum G, Tosco P, Kelley B, Ric, sriniker, gedeck, et al. rdkit/rdkit: 2022_09_1 (Q3 2022) Release 2022. 10.5281/zenodo.7235579.

[19] Xia H. Continuous-bag-of-words and Skip-gram for word vector training and text classification. J Phys Conf Ser 2023;2634:012052. 10.1088/1742-6596/2634/1/012052.

[20] Řehůřek R, Sojka P. Software Framework for Topic Modelling with Large Corpora. 2010. 10.13140/2.1.2393.1847.

[21] Wang N-N, Dong J, Deng Y-H, Zhu M-F, Wen M, Yao Z-J, et al. ADME Properties Evaluation in Drug Discovery: Prediction of Caco-2 Cell Permeability Using a Combination of NSGA-II and Boosting. J Chem Inf Model 2016;56:763–73. 10.1021/acs.jcim.5b00642.

[22] Hou T, Wang J, Zhang W, Xu X. ADME Evaluation in Drug Discovery. 7. Prediction of Oral Absorption by Correlation and Classification. J Chem Inf Model 2007;47:208–18. 10.1021/ci600343x.

[23] Broccatelli F, Carosati E, Neri A, Frosini M, Goracci L, Oprea TI, et al. A Novel Approach for Predicting P-Glycoprotein (ABCB1) Inhibition Using Molecular Interaction Fields. J Med Chem 2011;54:1740–51. 10.1021/jm101421d.

[24] Ma C-Y, Yang S-Y, Zhang H, Xiang M-L, Huang Q, Wei Y-Q. Prediction models of human plasma protein binding rate and oral bioavailability derived by using GA–CG– SVM method. J Pharm Biomed Anal 2008;47:677–82. 10.1016/j.jpba.2008.03.023.

[25] Wu Z, Ramsundar B, Feinberg EN, Gomes J, Geniesse C, Pappu AS, et al. MoleculeNet: a benchmark for molecular machine learning. Chem Sci 2018;9:513–30. 10.1039/C7SC02664A.

[26] Martins IF, Teixeira AL, Pinheiro L, Falcao AO. A Bayesian Approach to in Silico Blood-Brain Barrier Penetration Modeling. J Chem Inf Model 2012;52:1686–97. 10.1021/ci300124c.

[27] Document: Experimental in vitro DMPK and physicochemical data on a set of publicly disclosed compounds (CHEMBL3301361) n.d. https://www.ebi.ac.uk/explore/document/CHEMBL3301361 x(accessed July 3, 2025).

[28] Lombardo F, Jing Y. In Silico Prediction of Volume of Distribution in Humans. Extensive Data Set and the Exploration of Linear and Nonlinear Methods Coupled with Molecular Interaction Fields Descriptors. J Chem Inf Model 2016;56:2042–52. 10.1021/acs.jcim.6b00044.

[29] Veith H, Southall N, Huang R, James T, Fayne D, Artemenko N, et al. Comprehensive characterization of cytochrome P450 isozyme selectivity across chemical libraries. Nat Biotechnol 2009;27:1050–5. 10.1038/nbt.1581.

[30] Xu C, Cheng F, Chen L, Du Z, Li W, Liu G, et al. In silico Prediction of Chemical Ames Mutagenicity. J Chem Inf Model 2012;52:2840–7. 10.1021/ci300400a.

[31] Xu Y, Dai Z, Chen F, Gao S, Pei J, Lai L. Deep Learning for Drug-Induced Liver Injury. J Chem Inf Model 2015;55:2085–93. 10.1021/acs.jcim.5b00238.

[32] Karim A, Lee M, Balle T, Sattar A. CardioTox net: a robust predictor for hERG channel blockade based on deep learning meta-feature ensembles. J Cheminformatics 2021;13:60. 10.1186/s13321-021-00541-z.

[33] Popescu M-C, Balas VE, Perescu-Popescu L, Mastorakis N. Multilayer perceptron and neural networks. WSEAS Trans Cir Sys 2009;8:579–88.

[34] Akiba T, Sano S, Yanase T, Ohta T, Koyama M. Optuna: A Next-generation Hyperparameter Optimization Framework. Proc. 25th ACM SIGKDD Int. Conf. Knowl. Discov. Data Min., New York, NY, USA: Association for Computing Machinery; 2019, p. 2623–31. 10.1145/3292500.3330701.

[35] Watanabe S. Tree-Structured Parzen Estimator: Understanding Its Algorithm Components and Their Roles for Better Empirical Performance 2023. 10.48550/arXiv.2304.11127.

[36] Dubey SR, Singh SK, Chaudhuri BB. Activation functions in deep learning: A comprehensive survey and benchmark. Neurocomputing 2022;503:92–108. 10.1016/j.neucom.2022.06.111.

[37] Kingma DP, Ba J. Adam: A Method for Stochastic Optimization 2017. 10.48550/arXiv.1412.6980.

[38] [2310.00174] ADMET property prediction through combinations of molecular fingerprints n.d. https://arxiv.org/abs/2310.00174 (accessed July 3, 2025).

[39] KatanaGraph/SimGCN-TDC [Internet]. Katana Graph. Available: https://github.com/KatanaGraph/SimGCN-TDC. Accessed 2025 July 3.

