## Supplementary Table 1 for "Improving ADMET prediction with descriptor augmentation of Mol2Vec embeddings"

### **Supplementary Information**

[Roman Stratiichuk](https://orcid.org/0009-0001-6632-7642)^1,2^, [Nazar Shevchuk](https://orcid.org/0009-0003-8474-9376)^1^, [Roman Kyrylenko](https://orcid.org/0009-0009-4092-755X)^1^, [Volodymyr Vozniak](https://orcid.org/0009-0008-3055-5257)^7^, [Ihor Koleiev](https://orcid.org/0009-0001-4620-3175)^1,3^, [Taras Voitsitskyi](https://orcid.org/0000-0003-3127-3688)^1,3^, [Vladyslav Husak](https://orcid.org/0009-0000-7398-2330)^1,6^, [Zakhar Ostrovsky](https://orcid.org/0009-0003-4644-3587)^7^, [Ivan Khropachov](https://orcid.org/0009-0000-2025-2115)^1^, [Serhii Starosyla](https://orcid.org/0000-0002-5103-0635)^1^, [Semen Yesylevsky](https://orcid.org/0000-0002-6748-8931)^1,3,4,5^, [Alan Nafiev](https://orcid.org/0009-0004-8604-377X)^1^

^1^ Receptor.AI Inc., 20-22 Wenlock Road, London N1 7GU, United Kingdom.

^2^ Department of Biophysics and Medical Informatics, Educational and Scientific Centre “Institute of Biology and Medicine”, Taras Shevchenko Kyiv National University, 64 Volodymyrska Str., 01601, Kyiv, Ukraine.

^3^ Department of Physics of Biological Systems, Institute of Physics of The National Academy of Sciences of Ukraine, 46 Nauky Ave., 03038, Kyiv, Ukraine.

^4^ Institute of Organic Chemistry and Biochemistry, Czech Academy of Sciences, CZ-166 10 Prague 6, Czech Republic.

^5^ Department of Physical Chemistry, Faculty of Science, Palacký University Olomouc, 17. listopadu 12, 771 46 Olomouc, Czech Republic.

^6^ Department of Cellular, Computational and Integrative Biology, The University of Trento, Via Sommarive 9, 38123 Povo (Trento), Italy

^7^ Khmelnytskyi National University: Khmelnytskyi, Khmelnytskyi, UA

**Supplementary Table 1.** Descriptors used in Mol2Vec model with physico-chemical properties.

Molecular weight, logP, HBD, HBA, rotatable bonds, number of atoms, molar refractivity, TPSA, formal charge, heavy atoms, number of rings, number of aromatic rings, num valence electrons, num heteroatoms, fscp3, SAScore, Glaxo, Dundee, BMS, PAINS, SureChEMBL, MLSMR, Inpharmatica, LINT, Lipinski Rule of 5, Ghose Filter, Veber Filter, Rule of 3 Filter, REOS Filter, Drug-like Filter QED.

**Supplementary Table 2.** Tuned hyperparameters of the Mol2Vec-best model.

| **Dataset** | **Input features** | **Batch size** | **Learning rate** | **Weight decay** | **Activation** | **Number of layers** | **Number of units** | **Dropout values** | **Batch norm usage** |
| --- | --- | --- | --- | --- | --- | --- | --- | --- | --- |
| Caco2 | 1533 | 16 | 0.000654 | 0.000082 | ReLU | 4 | [3000, 2000, 4000, 2000] | [0.452087, 0.186705, 0.120004, 0.10092] | [True, False, True, True] |
| HIA | 663 | 128 | 0.000013 | 0.001456 | PReLU | 5 | [2000, 2000, 500, 500, 2000] | [None, None, 0.10078, 0.486959, 0.206879] | [False, False, True, True, True] |
| Pgp-inh | 1200 | 1024 | 0.005225 | 0.000062 | LeakyReLU | 1 | [1000] | [0.448155] | [True] |
| Bioavailability | 1683 | 2048 | 0.000015 | 0.000711 | ELU | 1 | [1000] | [0.220524] | [False] |
| BBB | 1841 | 64 | 0.000021 | 0.000128 | ReLU | 2 | [1000, 1000] | [0.293934, None] | [False, True] |
| PPBR | 2509 | 64 | 0.001358 | 0.000066 | ELU | 5 | [1000, 1000, 1000, 1000, 3000] | [0.37761, 0.289389, None, None, None] | [False, False, True, False, False] |
| VDss | 2125 | 64 | 0.000031 | 0.001702 | LeakyReLU | 5 | [4096, 2048, 8192, 1024, 2048] | [0.124507, None, None, 0.182864, None] | [True, False, True, False, True] |
| CYP2C9 Inhibition | 7577 | 2048 | 0.000012 | 0.001781 | ReLU | 2 | [4000, 4000] | [0.465949, None] | [True, True] |
| CYP2D6 Inhibition | 7307 | 512 | 0.000055 | 0.000435 | LeakyReLU | 2 | [8000, 4000] | [None, None] | [True, False] |
| CYP3A4 Inhibition | 8100 | 128 | 0.00001 | 0.000207 | LeakyReLU | 1 | [4000] | [0.423972] | [False] |
| CYP2C9 Substrate | 1776 | 256 | 0.011264 | 0.00013 | PReLU | 1 | [1000] | [None] | [False] |
| CYP2D6 Substrate | 1636 | 2048 | 0.007673 | 0.00012 | PReLU | 2 | [2000, 1000] | [None, None] | [False, True] |
| CYP3A4 Substrate | 2109 | 16 | 0.000077 | 0.000184 | PReLU | 1 | [2000] | [0.208099] | [False] |
| hERG | 1306 | 256 | 0.000022 | 0.003201 | PReLU | 1 | [1000] | [None] | [False] |
| AMES | 3642 | 256 | 0.000011 | 0.000448 | ReLU | 4 | [4000, 6000, 2000, 6000] | [0.480062, 0.145223, None, 0.221702] | [True, True, False, True] |
| DILI | 1151 | 1024 | 0.000011 | 0.002879 | ELU | 5 | [1000, 3000, 1000, 4000, 2000] | [0.422933, 0.124029, None, 0.121059, 0.115546] | [True, True, False, False, False] |

**Supplementary Table 3.** Comparison of four Mol2Vec-based featurization strategies - Mol2Vec Only, Mol2Vec with Physicochemical descriptors, Mol2Vec with Mordred descriptors, and Mol2Vec Best - across 16 TDC benchmarks. Arrows indicate whether higher (↑) or lower (↓) values are better.

| **Task** | **Metric** | **Mol2Vec Only** | **Mol2Vec PhysChem** | **Mol2Vec Mordered** | **Mol2Vec Best** |
| --- | --- | --- | --- | --- | --- |
| Caco-2 | MAE ↓ | 0.377 ± 0.029 | 0.367 ± 0.033 | 0.36 ± 0.031 | 0.315 ± 0.017 |
| HIA | ROC-AUC ↑ | 0.961 ± 0.018 | 0.968 ± 0.012 | 0.959 ± 0.021 | 0.996 ± 0.001 |
| Pgp-sub | ROC-AUC ↑ | 0.899 ± 0.008 | 0.896 ± 0.023 | 0.903 ± 0.005 | 0.948 ± 0.004 |
| Bioavailability | ROC-AUC ↑ | 0.633 ± 0.027 | 0.634 ± 0.059 | 0.736 ± 0.011 | 0.776 ± 0.027 |
| BBB | ROC-AUC ↑ | 0.916 ± 0.003 | 0.914 ± 0.007 | 0.923 ± 0.007 | 0.93 ± 0.004 |
| PPB | MAE ↓ | 10.964 ± 0.248 | 10.413 ± 0.146 | 10.931 ± 0.904 | 7.47 ± 0.192 |
| VD | Spearman ↑ | 0.312 ± 0.057 | 0.377 ± 0.088 | 0.425 ± 0.168 | 0.646 ± 0.027 |
| CYP2D6-inh | RR-AUC ↑ | 0.632 ± 0.016 | 0.639 ± 0.011 | 0.657 ± 0.006 | 0.726 ± 0.004 |
| CYP3A4-inh | RR-AUC ↑ | 0.837 ± 0.006 | 0.843 ± 0.007 | 0.851 ± 0.003 | 0.884 ± 0.001 |
| CYP2C9-inh | RR-AUC ↑ | 0.733 ± 0.011 | 0.741 ± 0.011 | 0.72 ± 0.02 | 0.8 ± 0.001 |
| CYP2D6-sub | RR-AUC ↑ | 0.678 ± 0.035 | 0.687 ± 0.022 | 0.636 ± 0.044 | 0.821 ± 0.008 |
| CYP3A4-sub | ROC-AUC ↑ | 0.598 ± 0.042 | 0.623 ± 0.022 | 0.608 ± 0.014 | 0.776 ± 0.015 |
| CYP2C9-sub | RR-AUC ↑ | 0.345 ± 0.033 | 0.353 ± 0.036 | 0.32 ± 0.026 | 0.556 ± 0.055 |
| hERG | ROC-AUC ↑ | 0.753 ± 0.03 | 0.781 ± 0.019 | 0.802 ± 0.022 | 0.897 ± 0.003 |
| AMES | ROC-AUC ↑ | 0.808 ± 0.004 | 0.823 ± 0.005 | 0.834 ± 0.003 | 0.876 ± 0.002 |
| DILI | ROC-AUC ↑ | 0.863 ± 0.034 | 0.861 ± 0.013 | 0.853 ± 0.019 | 0.964 ± 0.004 |
